# Spike conformational tuning rather than epitope erosion drives SARS-CoV-2 resistance to broadly neutralizing antibodies

**DOI:** 10.1101/2024.12.03.626570

**Authors:** Riku Fagerlund, Hasan Uğurlu, Petja Salminen, Eva Žusinaite, Anna R. Mäkelä, Tapio Kesti, Andres Merits, Ilona Rissanen, Kalle Saksela

## Abstract

Escape from protective immunity has been a defining feature of the COVID-19 pandemic and is widely attributed to mutations in neutralizing antibody epitopes within the receptor binding domain (RBD) of the SARS-CoV-2 spike glycoprotein. We investigated whether this paradigm accounts for the escape of Omicron lineage viruses from broadly neutralizing antibodies (bNAbs) targeting conserved RBD epitopes. We found that mutational erosion of bNAb binding epitopes was not a major driver of Omicron immune escape. Viruses carrying RBDs from highly resistant contemporary Omicron variants were potently neutralized by bNAbs when those RBDs were placed into an ancestral Wuhan-like spike backbone. Similarly, sera collected during the pre-Omicron era from individuals vaccinated with Wuhan-based vaccines retained robust RBD-specific neutralizing activity against these chimeric viruses. Mechanistically, bNAb treatment triggered exposure of the S2’ cleavage site in neutralization-sensitive but not resistant spike proteins, indicating that bNAb susceptibility reflects differences in spike propensity for antibody-induced premature conformational transition. These findings show that, rather than escaping through mutation of bNAb epitopes, SARS-CoV-2 has evolved spike variants with altered conformational dynamics that resist bNAb-induced destabilization of the prefusion spike trimer.

## INTRODUCTION

Rapid sequence evolution involving the neutralizing epitopes in the viral spike glycoprotein during the ongoing pandemic has reduced the ability of current vaccines to prevent infection and diminished the utility of approved therapeutic antibodies for treating COVID-19 [1]. The magnitude of this challenge escalated upon the emergence of the Omicron variant reported in South Africa in November 2021, and the subsequent spread of its subvariants [2-5].

Approximately 90% of all neutralizing antibodies (NAb) against SARS-CoV-2 target the receptor binding domain (RBD) of the spike protein [6, 7]. The majority of NAbs in COVID-19 patients and the first generation of therapeutic antibodies are specific for the receptor-binding motif (RBM) in the RBD, and are presumed to neutralize by preventing the RBD from engaging with the host receptor ACE2 [8]. However, RBM-binding antibodies are sensitive to SARS-CoV-2 sequence variation and tend to lose affinity against emerging variants [8].

By contrast, many NAbs that bind to conserved RBD regions outside of the RBM show broad neutralizing potency against divergent SARS-CoV-2 variants [9-16] as well as other sarbecoviruses, including SARS-CoV-1, and related animal viruses regarded as potential zoonotic threats. Such broadly neutralizing antibodies (bNAbs) typically target so-called cryptic sites in the RBD core region, which are occluded in the predominating closed conformation of the spike trimer where all or two out of the three RBDs are in “down-orientation” [8]. However, the inhibitory potency of these bNAbs is typically lower in comparison to RBM-targeting NAbs [8].

While the escape of emerging new SARS-CoV-2 variants from neutralization by RBM-targeting NAbs is driven by the accumulation of amino acid changes that alter their target epitopes, it remains less well understood how viral evolution may have contributed to the resistance of SARS-CoV-2 variants against bnAbs.

Here we report that, in comparison to the ancestral SARS-CoV-2 strain, most of the Omicron lineage variants show markedly increased bNAb resistance to pre-existing immunity. We show that in contrast to evasion from RBM-binding antibodies, escape from the cryptic site-targeting bNAbs does not appear to be driven primarily by target epitope loss but instead by altered conformational dynamics of the spike protein involving mutations outside of the RBD region.

## RESULTS

We examined the literature on bNAb clones obtained from COVID-19 patients and vaccinees to assemble a panel of structurally and functionally well-characterized antibodies. We did not include antibodies reported to have very low neutralization potencies and excluded bNAbs whose binding footprints substantially overlapped with the ACE2 binding interface. Accordingly, we chose to recombinantly express six prominent bNAbs, namely COVA1-16 [17], 10-28 [13], 10-40 [13], 2-36 [18], C022 [14], and S2H97 [19], five (COVA1-16, 10-28, 10-40, 2-36, C022) of which target the highly conserved class IV/CR3022 site [9] and one targeting the conserved site V (S2H97) in the RBD. A compilation of the structures of these antibodies in complex with RBD and survey of the associated binding footprints is provided in Figure 1.

**Figure 1.**
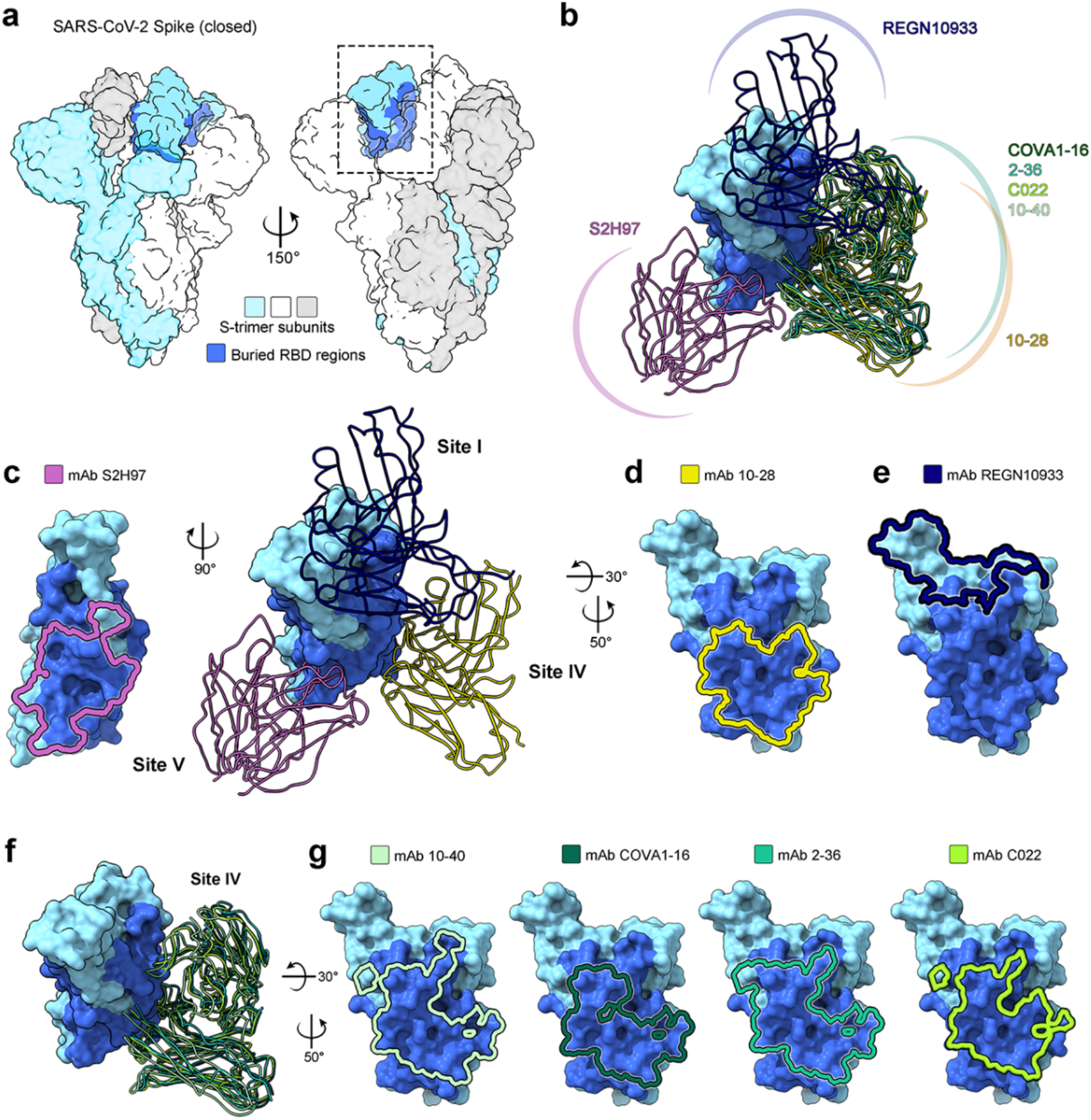
The epitopes targeted by the neutralizing antibodies (nAbs) used in this study. **A.** A surface representation of the SARS-CoV-2 S-trimer (PDB 6ZP0) [20]. The trimer subunits are shown in white, grey, and light blue, with the regions of the receptor-binding domain (RBD) that are inaccessible when the RBD is in the “down” or “closed” conformation highlighted in dark blue on one subunit. **B**. The antibodies examined in this study, namely COVA1-16 [17], 10-28 [13], 10-40 [13], 2-36 [18], C022 [14], and S2H97 [19], all bind regions of RBD that are buried in the down state. The epitope of mAb REGN10933 (Casirivimab) [21], which targets the ACE2-binding site, is shown for comparison. **C - E**. The binding modes and epitopes of mAb S2H97 (PDB 7M7W), 10-28 (PDB 7SI2) and REGN10933 (PDB 6XDG), which target antigenic sites V, IV and I, respectively. **F - G**. The binding modes and epitopes of mAbs COVA1-16 (PDB 7JMW), 10-40 (PDB 7SD5), 2-36 (PDB 7N5H), and C022 (PDB 7RKU), which target antigenic site IV. Epitope residues for each antibody-RBD complex were determined by the PDBePISA [22] tool for the exploration of macromolecular interfaces, and molecular graphics were generated with UCSF ChimeraX [23]. For clarity, only the Fv region of each antibody is shown, and all epitopes are shown mapped on the RBD from the S-trimer structure (PDB 6ZP0) [22].

The neutralization capacity of these antibodies was tested using a panel of pseudoviruses carrying spike glycoproteins of the prototypic B.1 virus (“Wuhan-D614G” in the following) or 10 diverse Omicron variants, including BA.1, BA.2, BA.4/5, BA.2.75 (Centaurus), XBB (Gryphon), BF.7 (Minotaur), BQ.1.1 (Cerberus), BU.1, and XBB.1.5 (Kraken), and EG.5.1 (Eris).

All six antibodies could efficiently neutralize Wuhan-D614G with IC_50_ values in agreement with the published data (Figure 2). By contrast, the Omicron variants were considerably more resistant to these antibodies, showing IC_50_ values typically more than one and, in many cases, more than two orders of magnitude higher than for Wuhan-D614G.

**Figure 2.**
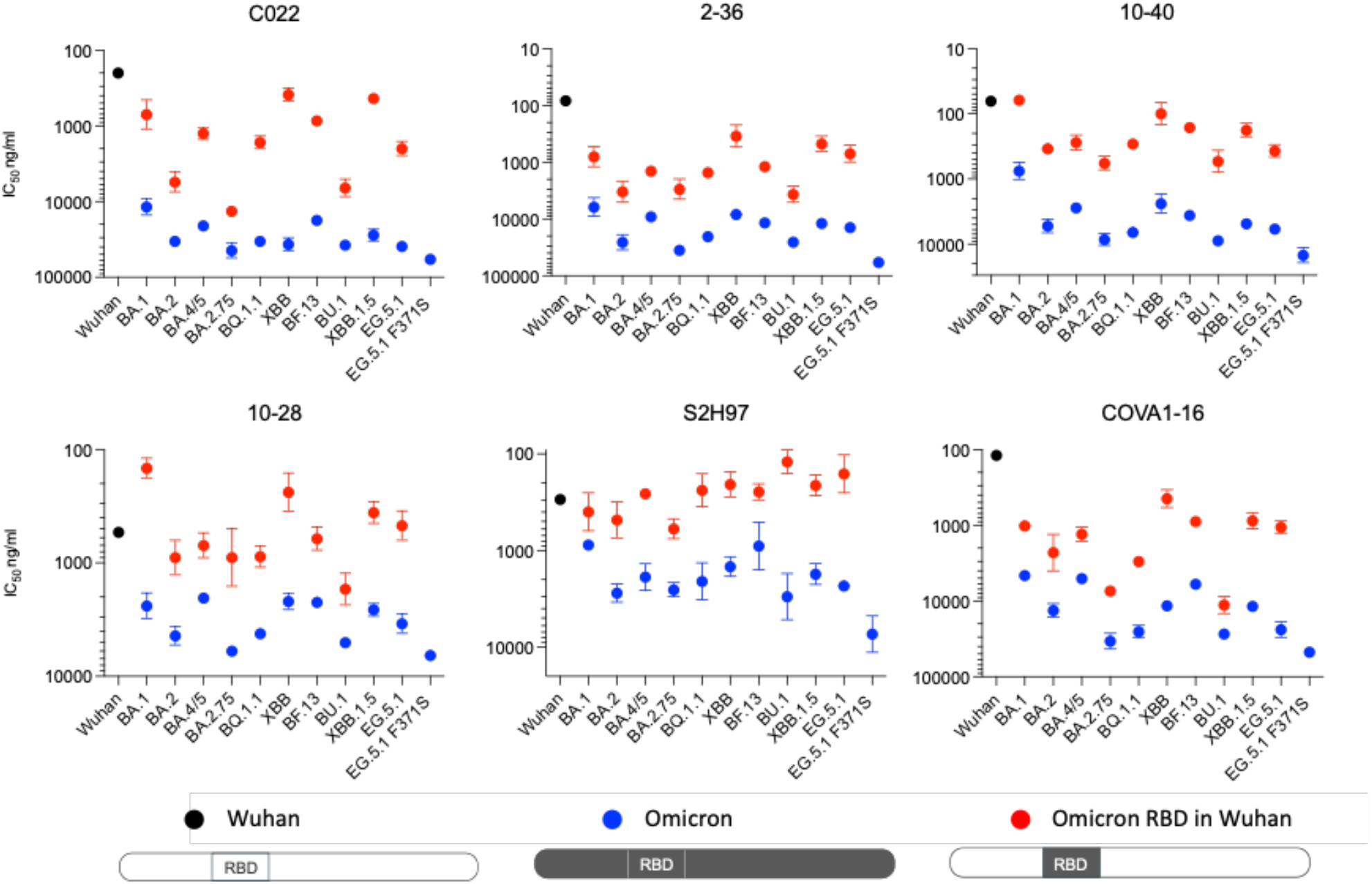
Neutralization by six prominent bNAbs of pseudoviruses carrying a spike protein from the ancestral Wuhan-D614G virus (black dot) or from 10 different Omicron variants (blue dots) or a chimeric spike containing the RBD (residues 330-529) of the same Omicron variants in the Wuhan-D614G backbone (red dots). Neutralization assays were repeated two or more times independently in duplicates. The averages and standard deviations of representative assays are shown. The original data are provided as a Source Data file.

Increased resistance against neutralization by RBD-binding antibodies is thought to be due their decreased capacity caused by RBD mutations that have accumulated during the immune escape-driven SARS-CoV-2 evolution [3, 24-30]. To further explore this concept, we constructed pseudoviruses carrying chimeric spike proteins containing the RBDs from the neutralization-resistant Omicron variants placed into the backbone of the Wuhan-D614G strain. Surprisingly, we observed that these heavily mutated Omicron RBDs did not transfer the resistant phenotype to Wuhan-D614G spike and, instead, the resulting chimeric spike proteins were remarkably easy to neutralize when compared to the corresponding native Omicron spike proteins (Figure 2). Of note, all of these neutralization-sensitive chimeric viruses carried a hydrophobic (L or F) substitution of RBD residue S371, which has been implicated as a cause for resistance against several classes of neutralizing antibodies [3, 28]. To further test the role of this residue, we introduced a F371S mutation into of EG.5.1 spike to restore a Wuhan-like serine residue in this position. This mutation showed no evidence of increased sensitivity of the EG.5.1 spike to neutralization by our panel of bNAbs (Figure 2).

These results prompted us to study sera from individuals immunized with the original Wuhan-based vaccines during the pre-Omicron era. In addition to XBB, BQ.1.1, XBB.1.5, and EG.5.1 used in the studies shown in Figure 2, another more recent Omicron variant JN.1 was included in this experiment. Five vaccine sera collected before February 2022 were tested and found to have strong neutralizing activity against the ancestral Wuhan-D614G virus (ID_50_ titers ranging between 5382 – 442; Figure 3). By contrast, in agreement with the well-documented immunoevasive capacity of these viruses [5, 29, 31-33], the neutralization potency of the vaccinee sera against these five Omicron variants was poor, showing titers that were more than an order of magnitude lower than observed for Wuhan-D614G and, in more than half of the cases, too low to be accurately determined. In agreement with the literature [6, 7], the neutralizing capacity of these sera was almost completely mediated by RBD-targeted antibodies, as 99% to 93% of this activity could be removed by immunodepletion with beads coated with recombinant RBD protein.

**Figure 3.**
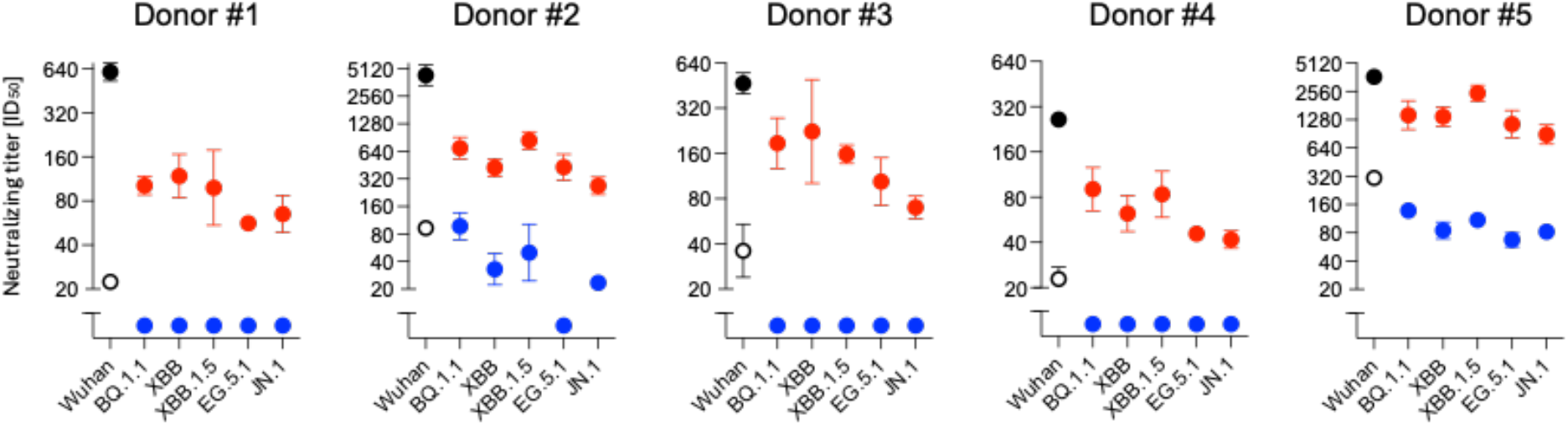
Neutralization of pseudoviruses carrying different spikes by human sera collected during the pre-Omicron era from persons immunized with original Wuhan-based vaccines. Blue dots indicate neutralization titres against viruses with the indicated native Omicron spikes, whereas red dots mark titres against chimeric viruses carrying the corresponding RBD (residues 330-529) transplanted into Wuhan-D614G backbone. Solid black dots show the titre against a virus with a native Wuhan-D614G spike, whereas empty black dots show the remaining neutralization titre after the same serum was immunodepleted with recombinant RBD protein. Neutralization assays were repeated twice in duplicates. The averages and standard deviations are shown. The original data are provided as a Source Data file.

In contrast to their poor potency against native Omicron viruses, all five vaccinee sera could effectively neutralize viruses in which the RBD of these Omicron variants was inserted into the Wuhan-D614G spike backbone, exhibiting ID_50_ titers that were only modestly decreased against the native Wuhan-D614G virus. Importantly, in all cases, the capacity to neutralize these chimeric viruses was much higher than the residual Wuhan-D614G-neutralizing activity of the same sera after immunodepletion with RBD protein, thereby excluding a significant contribution of Wuhan-D614G-specific non-RBD-targeting antibodies to neutralization by these sera.

Thus, we conclude that these human sera were rich in antibodies analogous to the six tested bNAb clones by potently targeting RBD epitopes that have remained intact during Omicron evolution but have lost their capacity for potent neutralization because of changes that have occurred elsewhere in the spike protein.

To explore the basis for the selective resistance of Omicron RBDs in their native context, we constructed reciprocal chimeras containing the Wuhan-D614G RBD placed in different Omicron spike backbones. This set of five Wuhan-D614G RBD-containing chimeric spikes covers nine of the ten Omicron variants tested in Figure 1, as four of them share the same backbone and differ only in their RBD sequences. When tested against the panel of six bNAbs, the neutralization sensitivity of these reciprocal spike chimeras did not generally differ from Wuhan-D614G (Figure 4).

**Figure 4.**
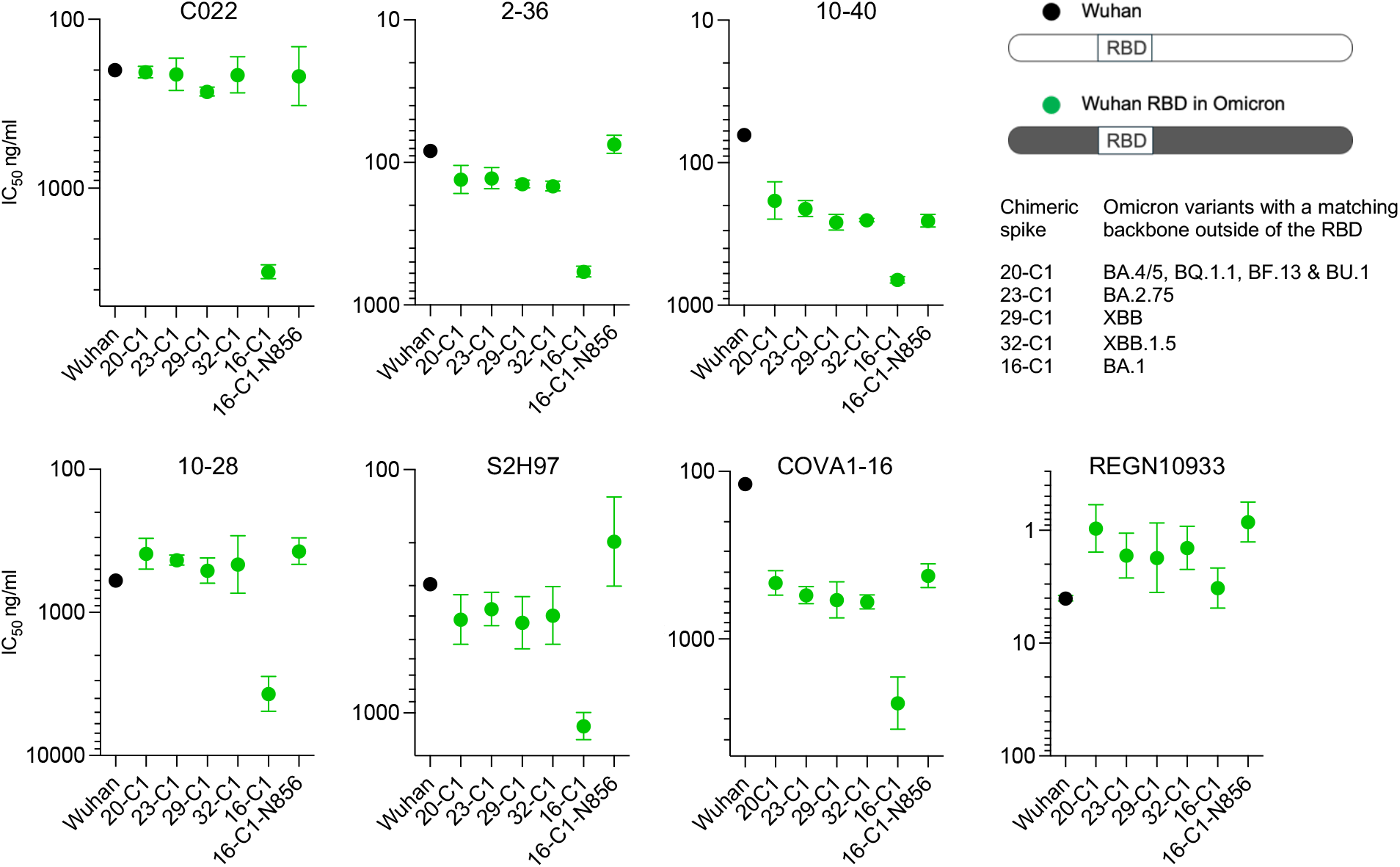
Neutralization of pseudoviruses with chimeric spike proteins by different antibodies. Round symbols indicate neutralization (half-maximal inhibitory mAb concentration) of pseudoviruses containing Wuhan-D614G spike (black dots) or chimeric Omicron spikes containing Wuhan-D614G RBD (green dots). The Omicron backbone of 20-C1 corresponds to the spikes of BA.4/5, BQ.1.1, BF13, and BU.1, whereas 23-C1 is BA.2.75, 29-C1 is XBB, 32-C1 is XBB.1.5, and 16-C1 is BA.1. Data are also shown using a virus pseudotyped with a 16-C1 spike containing a K-to-N back-mutation restoring the Wuhan-like asparagine at position 856. Neutralization assays were repeated independently two or more times in triplicates. The averages and standard deviations of representative assays are shown. The original data are provided in a Source Data file.

Together with our earlier results, this finding indicated that the resistant phenotype of the Omicron spike proteins cannot be simply assigned to their RBD domain nor to the remaining spike backbone alone, but instead appears to be determined by the intramolecular dynamics of the entire spike protein.

An interesting exception to the general neutralization sensitivity of the Wuhan-D614G RBD-containing Omicron spike proteins was the chimera 16-C1 composed of the BA.1 spike backbone, which was markedly difficult to neutralize by these bNAbs. As a control antibody that does not target a cryptic RBD core epitope, we tested REGN10933 (Casirivimab), a canonical ACE2-competing RBM-binding antibody [21], which efficiently neutralized also the 16-C1 chimeric spike virus.

This unusual behavior of 16-C1 led us to note that BA.1 spike contains a set of unique mutations in its S2 region that are lacking in subsequent Omicron variants. One of these, namely N856K, is located C-terminally adjacent to the fusion peptide at an interprotomer S1/S2 interface where it could potentially influence molecular interactions within the spike trimer. We therefore tested the effect of introducing the original Wuhan-like asparagine residue into position 856 of the 16-C1 spike. Remarkably, we found that this single back-mutation effectively abolished the bNAb-resistance of 16-C1 (Figure 4).

To investigate the mechanism by which bNAbs inhibit spike-mediated infectivity, and how Omicron variants acquire resistance to this inhibition, we examined the exposure of the S2’ protease cleavage site in spike trimers on the surface of virions incubated in the presence or absence of bNAbs (Figure 5). Normally the S2’ site is exposed upon ACE2 engagement, which promotes cleavage of a 13 kDa N-terminal fragment from the S2 subunit by membrane-associated TMPRSS2 or by endosomal cathepsins, thereby allowing the spike to complete the transition of the spike trimer from the prefusion to the postfusion conformation (Figure 5D).

**Figure 5.**
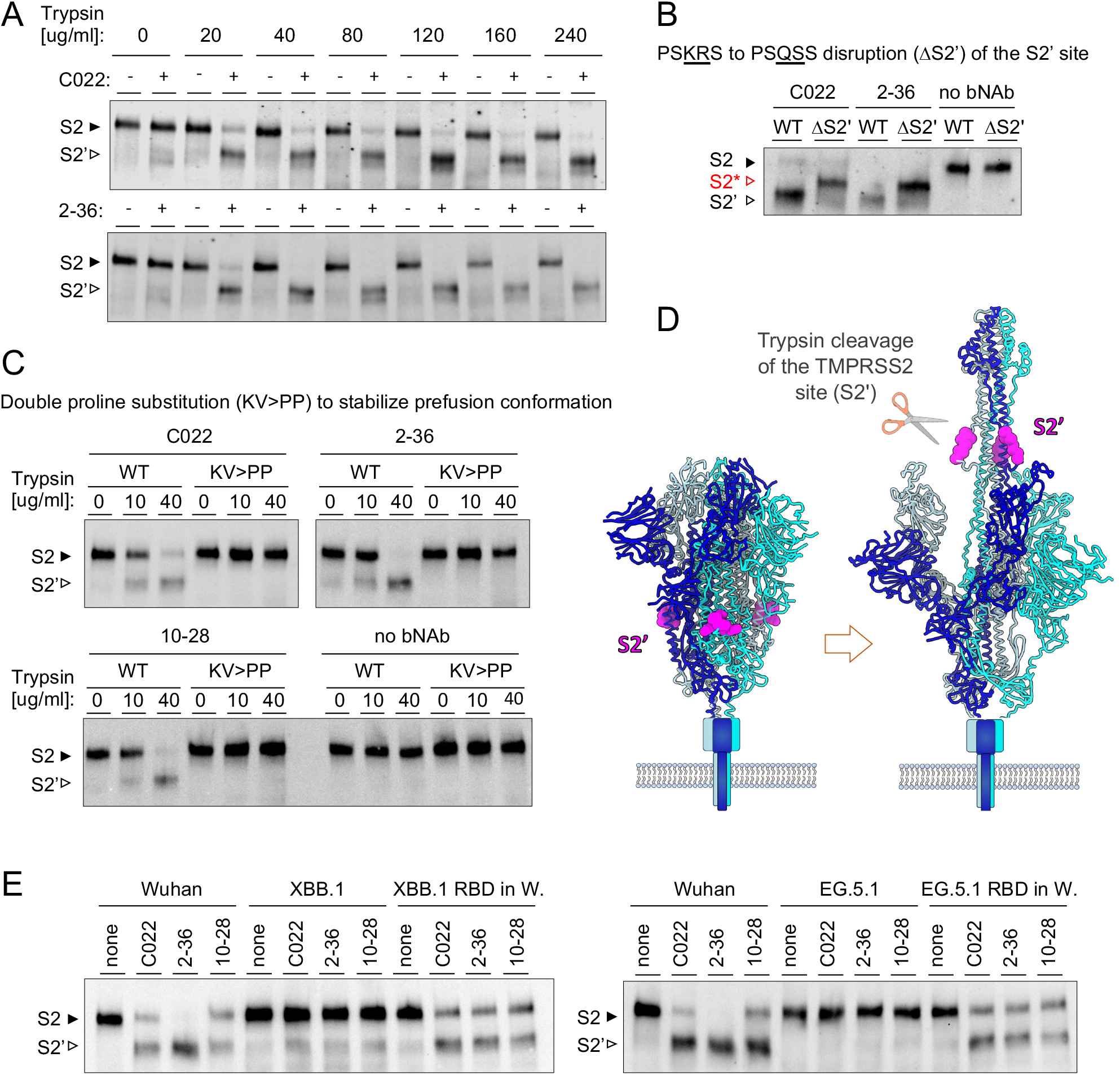
Exposure of S2’ site by bNAbs. **A.** Virions having Wuhan-D614G spike were incubated with or without C022 or 2-36 and treated with the indicated concentrations of trypsin. **B**. Virions having Wuhan-D614G (WT) spike or a mutant of it with a disrupted TMPRSS2 site (ΔS2’) were incubated with C022, 2-36, or without an antibody (no bNAb) and followed by treatment with 40 µg/ml of trypsin. **C**. Virions having Wuhan-D614G spike (WT) or prefusion-stabilized K986P/V987P mutant of it (KV>PP) were incubated with C022, 2-36, or 10-28 or without an antibody and treated with the indicated consentrations of trypsin. **D**. Ilustrustration of the structural basis of the TMPRSS2/trypsin cleavage (S2’) site exposure upon spike trimer transition from its prefusion conformation to an early fusion intermediate drawn according to the cryo-EM structures PDB 6VXX [35] and PDB 8Z7P [36]. **E**. Virions with Wuhan-D614G, XBB, EG.5.1, or chimeric spikes containing XBB-RBD or EG.5.1-RBD in the Wuhan backbone were incubated with C022, 2-36, 10-28, or without the antibody and treated with 40 µg/ml of trypsin. All membranes were stained with an anti-S2 antibody.

Using the bNAbs C022 and 2-36, we observed that antibody treatment strongly sensitized virion-associated Wuhan-D614G spike protein to proteolytic cleavage at the S2’ site by trypsin, a soluble protease with TMPRSS2-like catalytic mechanism and substrate specificity [34] (Figure 5A). When a spike mutant with a disrupted S2’ site was examined, bNAb treatment still sensitized the spike to trypsin digestion, but the altered molecular weight of the S2 cleavage product(s) indicated that the bNAb-exposed trypsin cleavage site in the native spike is the authentic S2’ site (Figure 5B). By contrast, when virions carrying a prefusion-stabilized K986P/V987P spike were examined, these antibodies no longer promoted S2’ cleavage, confirming that enhanced proteolytic susceptibility results from bNAb-induced destabilization of the prefusion conformation (Figure 5C).

Having validated this experimental approach, we next examined whether spike proteins from the Omicron variants XBB and EG.5.1 are similarly susceptible to bNAb-mediated sensitization. To address this, we tested three bNAbs (C022, 2-36, and 10-28). In marked contrast to virions bearing Wuhan-D614G spike or chimeric spikes containing only the XBB or EG.5.1 RBDs, virions carrying native XBB or EG.5.1 spike trimers showed little sensitization to S2’ cleavage following bNAb treatment (Figure 5E).

Together, these findings provide direct biochemical evidence that neutralization by these bNAbs is mediated by destabilization of the spike prefusion conformation, whereas resistance of the tested Omicron variants involves stabilization of the infection-competent prefusion state.

## DISCUSSION

Our findings provide new mechanistic insight into SARS-CoV-2 immune evasion, demonstrating that altered spike protein dynamics enable Omicron variants to escape broadly neutralizing human antibodies (bNAbs) without altering the highly conserved RBD residues that form their binding epitopes.

Whereas the conservation of cryptic RBD epitopes targeted by several bNAbs is well recognized, the reasons for their progressive loss of neutralization potency against Omicron variants has not received much attention. This may be partly because, in comparison to many ACE2 interface-targeting antibodies, the most archetypal bNAbs like CR3022 [9] show relatively modest neutralizing activity even against ancestral SARS-CoV-2 strains.

It is therefore important to point out that a very substantial fraction of the neutralizing activity in sera from individuals immunized in the pre-Omicron era with a Wuhan-based vaccine is directed against epitopes that remain conserved in contemporary Omicron variants, as demonstrated by our RBD-swapping experiments. Nevertheless, the magnitude of the conformational escape from this immunity was striking. Although the Omicron variant EG.5.1 exhibited near-complete resistance to neutralization by vaccinee sera, transplantation of the EG.5.1 RBD into an ancestral spike backbone restored susceptibility, allowing RBD-targeted antibodies in these sera to neutralize the chimeric virus almost as efficiently as a virus bearing the Wuhan-D614G spike.

Notably, most of the SARS-CoV-2 neutralizing activity in these vaccinee sera was directed against the RBD and could be efficiently depleted using recombinant RBD. Therefore, the potent neutralization of viruses carrying chimeric spikes containing Omicron RBDs was not attributable to Wuhan-specific, non-RBD-targeting antibodies. Interestingly, a previous study by Bukh and colleagues showed that a panel of chimeric SARS-CoV-2 viruses generated by reverse genetics, in which RBDs from eight contemporary Omicron variants were inserted into the backbone of the Wuhan-like ancestral DK-AHH1 isolate, were neutralized by sera from vaccinated and convalescent individuals much more efficiently than the corresponding viruses carrying full-length native spikes from the same Omicron variants [37]. This observation is in good agreement with our current data. However, Bukh et al. did not assess the contribution of RBD-targeted antibodies in these sera. Instead, they speculated that neutralizing antibodies directed against the N-terminal domain, capable of binding to DK-AHH1 but not to Omicron spike proteins, might explain their findings.

We demonstrate that the bNAbs studied here neutralize SARS-CoV-2 by displacing the spike trimer from its prefusion conformation, thereby exposing the TMPRSS2 cleavage site and inducing a state that likely corresponds to the early fusion intermediate recently described by Xing et al. [36]. A key observation of our study is that susceptibility of virion-associated spike proteins to this conformational transition determined the sensitivity to bNAb-mediated neutralization.

The idea that antibodies targeting cryptic RBD epitopes may neutralize by destabilizing the spike is not new. However, previous studies relied on recombinant spike ectodomains that may not faithfully recapitulate native spike dynamics, particularly as these constructs were engineered with K986P/V987P or hexaproline substitutions to enforce their prefusion state. Our findings not only substantiate the model of bNAb-mediated neutralization through prefusion destabilization in the context of native, virion-associated spike, but also provide important insight into the more elusive question of how Omicron variants have evolved resistance to these antibodies.

One proposed explanation has been that mutations within the Omicron RBD stabilize the down-conformation and thereby limit exposure of cryptic epitopes that are optimally accessible only when all three RBDs are in the up conformation (3-RBD-up). Specifically structural analyses of BA.1 and BA.2 spike proteins indicated that substitution of the RBD residue S371 with hydrophobic amino acids (L or F in all Omicron spikes) promotes the 3-RBD-down and 1-RBD-up states of [38-41], whereas introduction of the S371L/F substitution into the Wuhan spike confers substantial resistance to several bNAbs [3, 28, 42]. However, we found that despite containing the S371L/F mutation, all chimeric spike proteins carrying various Omicron RBDs remained readily neutralized by the tested bNAbs. Cryo-EM structures suggest that the hydrophobic substitution at position 371 limits RBD-up conformations by stabilizing interprotomer RBD interactions within Omicron spike trimers. Although such RBD–RBD interactions would be expected to persist in our Omicron RBD-containing chimeric spikes, they were insufficient to confer bNAb resistance. Conversely, reverting residue 371 back to serine failed to render the native EG.5.1 spike susceptible to neutralization. Thus, although the S371L/F substitution may contribute to stabilization of RBD-down states when present together with other naturally accumulated Omicron RBD mutations, it is neither necessary nor sufficient for resistance to these bNAbs.

Regardless of the precise contribution of the residue 371, our findings indicate a role for broader changes in spike conformational dynamics beyond just altered RBD up/down propensity, and demonstrate that amino acid changes in the Omicron spike outside the RBD are essential for bNAb resistance. However, as shown by our reverse chimera experiments (Omicron spikes containing the Wuhan RBD), mutations in the non-RBD spike framework are not sufficient to confer bNAb resistance on their own and require accompanying changes within the Omicron RBDs. These interacting networks of RBD and non-RBD mutations remain to be determined and are likely to differ between individual Omicron variants. Some insight into this question was provided by the striking bNAb resistance of the chimeric spike protein 16-C1, in which the ancestral Wuhan-D614G RBD was introduced into the BA.1 backbone. Restoration of a single lysine residue in the S2 region to the Wuhan-like asparagine (K856N) was sufficient to render 16-C1 readily neutralizable. Analysis of available SARS-CoV-2 spike structures in different conformational states shows that residue 856 is located distal to the RBD-spike interface (Figure S1), in a position where it may allosterically modulate interprotomeric S1/S2 interactions. Thus, N856K substitution could hinder spike trimer opening and thereby reduce susceptibility of the 16-C1 prefusion spike to bNAb-induced destabilization.

Collectively our findings underscore the versatility with which viruses adapt to humoral immune pressure. When neutralizing epitopes lie within conserved and functionally constrained regions of an envelope protein, escape can instead arise through mutations at distal sites that reshape the overall conformational landscape rather than through direct alteration of the epitope, which may impose a substantial fitness cost.

From a translational perspective these observations raise the interesting possibility that small-molecule antivirals designed to promote a similar conformational transition in the SARS-CoV-2 spike trimer could represent a novel therapeutic strategy. If such compounds could be developed to be less susceptible than antibodies to the conformational resistance mechanisms that have emerged in Omicron variants, they might act synergistically with existing bNAbs and help unleash the substantial yet largely latent neutralizing potential present in SARS-CoV-2–seropositive individuals, as revealed in this study.

## MATERIALS AND METHODS

### Antibody production

HEK293T cells (ATCC; Catalogue number CRL-3216) were maintained in high-glucose Dulbecco’s Modified Eagle Medium (Sigma Aldrich) supplemented with 10% fetal bovine serum, 1% L-Glutamine, and 1% penicillin/streptomycin (complete cell culture medium). The genes encoding the heavy and light chains variable domains of COVA1-16 [17] (7JMX_H and 7JMX_L), 10-28 (7SI2_H and 7SI2_L) [13], 10-40 (7SD5_H and 7SD5_L) [13], 2-36 (7N5H_J and 7N5H_K) [18], C022 (7RKU_J and 7RKU_K) [14], and S2H97 (7M7W_C and 7M7W_D) [19] were ordered as synthetic codon-optimized cDNA fragments (Integrated DNA Technologies), and seamlessly fused using GeneArt Gibson assembly cloning kit (Invitrogen) to the corresponding constant regions (from IgG1 heavy chain allotype IGHG1*01, in the mammalian expression vector pEBB [Addgene 22226]). The heavy and light chain containing vectors were co-transfected into HEK293T cells in complete cell culture medium supplemented with ultra-low IgG FBS (Gibco) using TransIT-2020 reagent (Mirus Bio) according to the manufacturer’s instructions. Supernatants containing the secreted human monoclonal antibodies were collected 48-96 h post-transfection, filtered through a 0.2 µm polyethersulfone membrane (Fisherbrand; Catalogue number 15913307), and concentrated using centrifugal filters (Amicon Ultra; catalogue number UFC901008). The final human IgG concentrations were determined using an in house anti-human IgG sandwich-ELISA.

### Pseudoviruses

HEK293T cells grown in T175 flask were transfected by using TransIT-2020 reagent (Mirus Bio) with the packaging plasmid p8.9NDSB (Addgene #132929), genomic plasmid pWPI (Addgene # 12254) expressing Renilla luciferase, and the expression vector pEBB (Addgene 22226) containing different codon-optimized SARS-CoV-2 spike glycoprotein cDNAs in which the last 18 codons were deleted to enhance the plasma membrane transport. The amino acids sequences for the spike variants used were based on consensus sequences for different SARS-CoV-2 variants according to information from the GISAID initiative (https://gisaid.org) curated by the outbreak.info project [43]. The culture media was replaced 12–16 h post-transfection with complete cell culture medium. The supernatant containing the SARS-CoV-2 spike glycoprotein-harboring pseudoviruses was collected 48 h after transfection and passed through a 0.22µm filter.

### Neutralization assays

Pseudovirus neutralization assays were performed essentially as previously described [44]. Briefly, serial dilutions of bNAbs were incubated with SARS-CoV-2 spike pseudotyped lentiviral vectors, after which ACE2-HEK293T cells [44] maintained in complete cell culture medium were added. Productive infection was quantified using Renilla-GLO assay (Promega; catalogue number E2710) and measured using the GloMax Navigator luminometer (Promega, catalogue number GM2010).

Half maximal inhibitory concentrations (IC_50_) and half maximal inhibitory dilutions (ID_50_) were determined using Prism10 software (GraphPad). To increase the accuracy of curve fitting, the control value (no antibody added) was set to the average of the measured control value and the values above it in such dilution series where luciferase readings from more than one of wells at the lowest antibody concentrations were higher than in the no-antibody control well.

The human sera used for neutralization assays were collected between September 2021 - January 2022 under informed consent from persons who had received 2 or 3 doses of the BNT162b2 mRNA vaccine with no history of SARS-CoV-2 infection.

### Trypsin cleavage

Spike pseudotyped virions were incubated in the absence or presence of bNAbs at 37°C for 3 hours. Reactions were cleaned with Capto™ Core 700 according to the manufacturer’s instructions. Cleaned reactions were treated with trypsin (Cytiva HyClone™) at 37°C for 15 minutes before the reaction was stopped by adding 4x Laemmli sample buffer containing beta-mercaptoethanol. Western blot analysis was performed according to standard protocols.

### Molecular graphics

Molecular graphics and analyses were performed with UCSF ChimeraX, developed by the Resource for Biocomputing, Visualization, and Informatics at the University of California, San Francisco, with support from National Institutes of Health R01-GM129325 and the Office of Cyber Infrastructure and Computational Biology, National Institute of Allergy and Infectious Diseases [23].

## ACKNOWLEDGEMENTS

We thank Virpi Syvälahti and Sanna Mäki for expert technical assistance. We gratefully acknowledge Ian A. Wilson and Meng Yuan for their helpful comments on an earlier version of this work.

## DATA AVAILABILITY

All data generated or analysed during this study are included in the main text and its supplementary information files.

## SUPPLEMENTARY DATA

**Figure S1.**
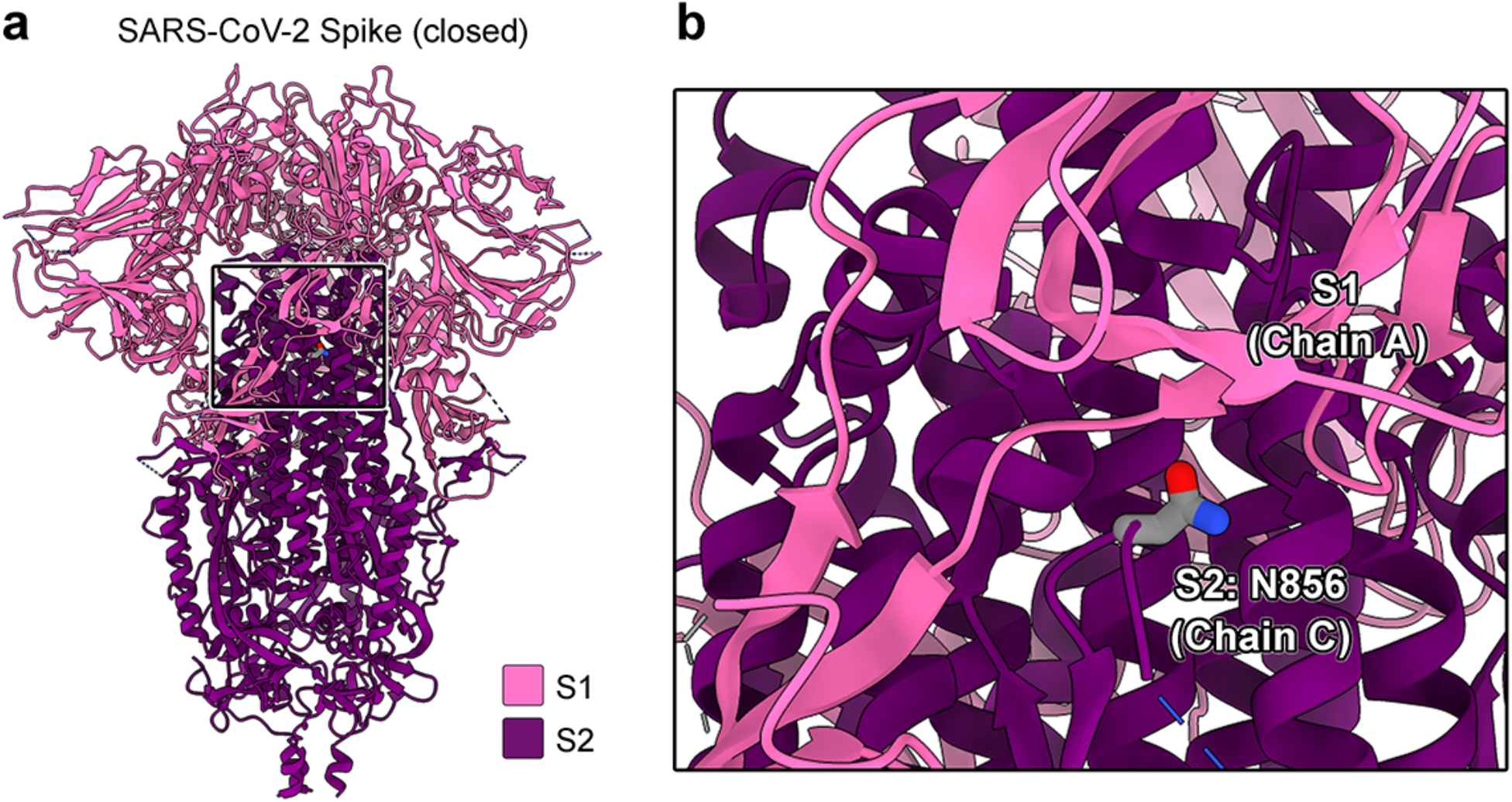
Residue N856 (K856 in BA.1) within SARS-CoV-2 S-trimer S2 is located at an inter-subunit interface. **A**. A cartoon representation of the SARS-CoV-2 S-trimer (PDB 6ZP0) [20]. The trimer subunits are colored pink for the S1 region (residues up to 667) and dark purple for the S2 region (residues 668 and beyond). **B**. A close-up panel displaying N856 in stick representation. Molecular graphics were generated with UCSF ChimeraX [23].

